# Single-cell Transcriptomics Analyses Revealed Specialized Microglial Subsets with Oligodendrocyte-like Signatures

**DOI:** 10.64898/2026.05.11.724239

**Authors:** Yanting Luo, Xuemei Huang, Yadan Nie, Haixia Wang, Zuoli Sun, Jian Yang, Yi He

## Abstract

**Background:** Microglial heterogeneity is a fundamental feature of brain homeostasis and pathology. The purpose of this study was to investigate the complexity of microglial plasticity by characterizing specialized oligodendrocyte-like microglial subsets.

**Methods:** The study was performed utilizing single-cell transcriptomics analyses and immunofluorescence staining to identify and profile microglial subpopulations. Additionally, spatial transferring and morphological analyses were conducted to determine the anatomical distribution and structural features of these specific cells.

**Results:** We identified a distinct microglial subset termed dual-phenotype microglia (DPM), which co-expresses microglial and oligodendrocyte markers. DPM consisted of two subtypes with distinct functions: myelin-associated DPM (mDPM) and neuron-associated DPM (nDPM). Spatial and morphological evaluations revealed that mDPMs were sparsely distributed across the whole brain and exhibited a highly ramified architecture, whereas nDPMs were enriched in the hippocampal dentate gyrus. Mechanistically, we found that mDPM function was driven by the Sox10 regulon to modulate myelin maintenance and axonal ensheathment, while nDPM was orchestrated by Glis2, facilitating essential neuron-glia crosstalk and synaptic regulation. Furthermore, we demonstrated that nDPM and mDPM were predicted to undergo significant alterations in multiple sclerosis and Alzheimer’s disease. Notably, mDPMs were selectively enriched in active multiple sclerosis lesions, revealing that DPM were closely related to neuropsychiatric disorders.

**Conclusions:** By comprehensively characterizing the morphology, molecular signatures, and spatial logic of these oligodendrocyte-like microglial subsets, our study elucidated the complexity of microglial plasticity. These findings provided new insights into their diverse roles in central nervous system health and disease.

**Graphical abstract:** Identification, Molecular Profiling, and Functional Modeling of Dual-Phenotype Microglia (DPM). (**1**) Discovery: Identification of the dual-phenotype microglia (DPM) population through single-cell transcriptomics. (**2**) Molecular Signatures: The transcriptomic identity of DPM subtypes is governed by specific regulatory networks. (**3**) Distribution & Pathology: Spatial mapping reveals divergent anatomical logic and disease relations for DPM subtypes. (**4**) Mechanism/Theory: A proposed functional model of mDPMs as “metabolic relay” and support units.

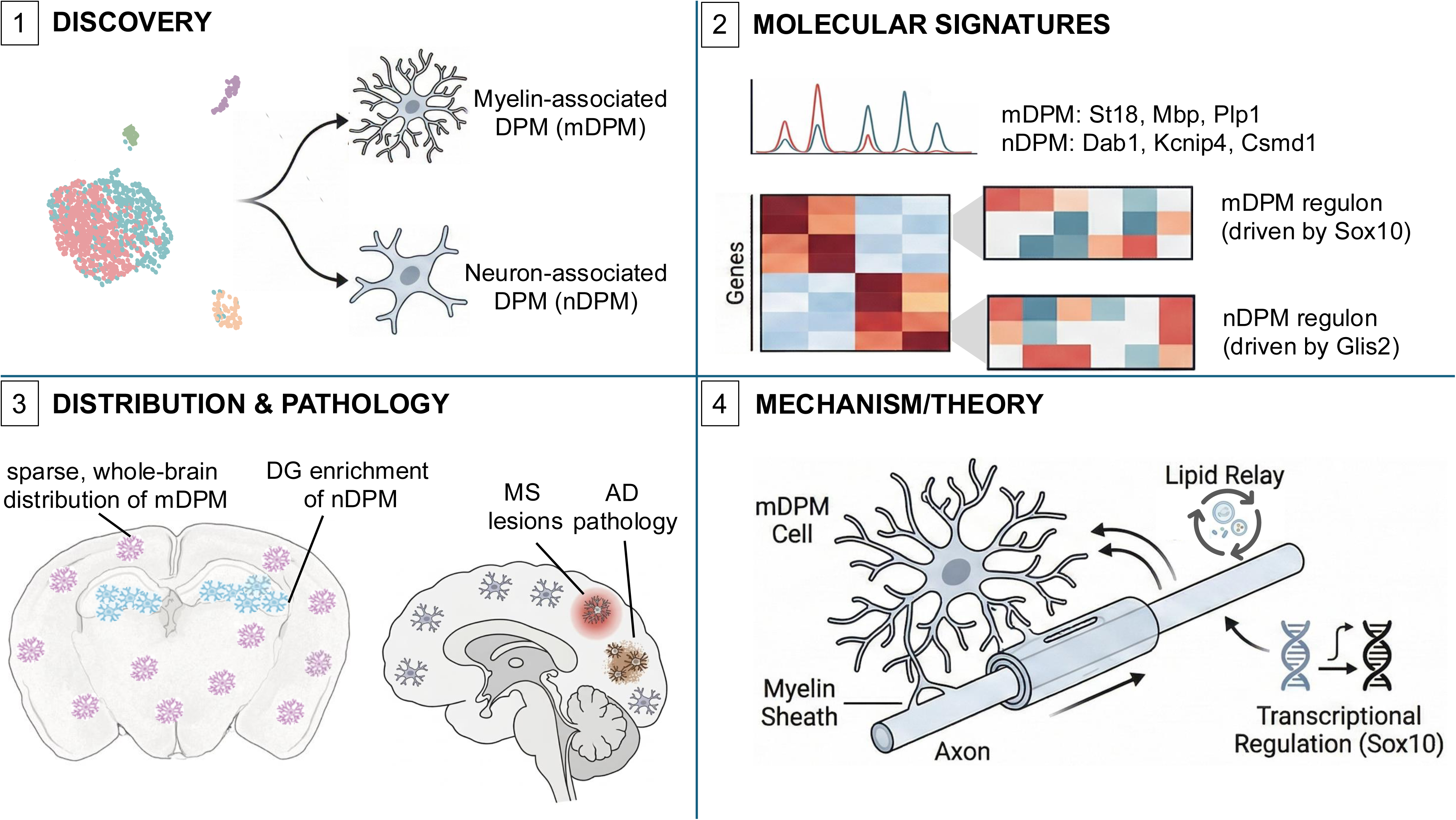

## Background

Microglia characterized highly heterogeneous cell population with the gene expression profiles dynamically vary with age, brain region, and microenvironment. Their morphological, ultrastructural, and molecular characteristics demonstrate considerable plasticity, leading to the coexistence of distinct cellular states that are closely linked to diverse functional roles(1). Recent advances in single-cell and single-nucleus RNA sequencing (scRNA-seq and snRNA-seq) have further elucidated these dynamic phenotypic changes over time and across neuroanatomical regions(2). And some studies have unveiled a spectrum of diverse microglial cell states in both healthy and diseased brains (3). With brain aging and in age-related neurodegenerative conditions, microglial phenotypes undergo regional and temporal evolution. Beyond their unique transcriptional profiles, reactive microglial subtypes such as ARM, DAM, MGnD, LDAM, and WAM share fundamental impairments in lipid metabolism and phagocytic capacity. These overlapping traits suggest that the functional identity of these cells is intimately coupled with their underlying metabolic state(4–6).

Beyond the classical roles in phagocytosis and pro-inflammatory responses, microglia are now known to shape myelination processes, an aspect that has drawn increasing attention. During neurodevelopment, activated microglia in the subventricular zone secrete cytokines such as TNFα, IL-1β, IL-6, and IFN-γ, thereby promoting oligodendrocyte development. Reductions in these cytokine levels have been shown to impair oligodendrocyte and consequently disrupt myelin formation(Shigemoto-Mogami et al., 2014). An early study performing microglia-oligodendrocyte co-culture indicated that microglia can activate oligodendrocytes to synthesize the myelin-specific galactolipid sulfatide, as well as myelin proteins such as *MBP* and *PLP* (7). While previous researches focused on microglia-oligodendrocyte interactions in the context of myelination, recent studies suggested that some microglia may intrinsically exhibit oligodendrocyte-like attributes. For instance, Liang C *et al.* identified a transcriptional cluster co-expressing microglial and oligodendrocytic genes significantly, annotating it as “Oligodendrocyte-Microglia”(8). Su Y *et al.* reported a microglial subset associated with “myelination” and “axon ensheathment” — classical gene functions of oligodendrocytes(9). However, these studies have not yet deeply characterized the localization and defining features of this particular cell population.

In our study, we identified a previously uncharacterized cell population that concurrently exhibits both microglial and oligodendrocytic transcriptional traits. We have termed these cells Dual-Phenotype Microglia (DPM). Building upon our previously published single-cell transcriptomic data, we investigate the DPM cells comprehensively, including functional characteristics and spatial distribution within the brain, and further explore their potential role in pathological contexts. Our work aims to shed light on this intriguing cellular phenotype, offering new insights into its biological and clinical significance.

## Methods

### Animals

Mice C57Bl/6J were used in this study. All animals were maintained in a specific pathogen-free facility with food and water ad libitum and maintained on a 12-h light/dark cycle.

### Immunofluorescence

Mice were anesthetized (intraperitoneal 50 mg of 1% Pelltobarbitalum Natricum per kilogram of body weight) and transcardially perfused with PBS. Brains were dissected and postfixed for 10-12 hours at 4°C in 4% paraformaldehyde, washed with PBS and gradient dehydrated in sucrose at 4°C. Samples were embedded and frozen in O.C.T. (Tissue-Tek). For all experiments, 40-μm coronal brain sections were obtained with a cryostat (Leica). The mounted brain sections were washed 3 times with PBS before permeating by 0.3% Triton X-100 in PBS for 15 min and blocking with 10% donkey serum in 0.1% Triton X-100 for 2h at room temperature. Then, sections were incubated with primary antibodies (IBA1, 1:1000, Abcam, Cat. ab5076; ST18, 1:100, Thermofisher, Cat. PA5-40764; DAB1, 1:100, Thermofisher, Cat. PA5-62538;) overnight at 4°C. Brain sections were incubated with specific fluorophore-labeled secondary antibodies (Alexa fluor 576, Alexa fluor 488, Alexa fluor 647, Thermofisher) for 1 hour at room temperature after washing 3 times for 10 min each in PBS. Then, brain sections were mounted by antifade mounting medium with DAPI (Solarbio, Cat. C0065) following 3 times wash. Images were obtained with a confocal microscope (Leica STELLARIS 5).

Hippocampal images were acquired from brain sections using z-stack scanning at 40× magnification, with a step size of 0.3 μm for three-dimensional reconstruction. A total of 3 mice were used, and 6–8 fields of view were captured per hippocampal hemisphere. Three-dimensional (3D) digital reconstruction of microglial morphology was performed using the Filament Tracer module (Imaris 10.1.0). The Autopath algorithm was employed to trace microglial processes based on fluorescence intensity and local contrast. For 2D analysis, z-stacks were converted into Maximum Intensity Projections (MIP) using Fiji. After binarization and skeletonization of the Iba1 signal, Sholl analysis was performed using the Fiji plugin (Neuroanatomy), with the starting radius set at 5 μm and a radius step size of 5 μm from the center of the cell soma (10).

### Cell clustering and visualization

The single-cell RNA sequencing data analyzed in this study were derived from our previously published dataset of hippocampal cells (11). The original dataset comprised 64,399 single-cell transcriptomes obtained from three wild-type (WT) and three IL15RA knockout (KO) mice. For this study, we performed a re-analysis specifically on the microglial cell subset isolated from the three WT mice. Microglial cells were identified based on canonical marker genes (e.g., *Cx3cr1* and *P2ry12*). Following standard preprocessing, the raw gene expression matrix was normalized using the LogNormalize method and subsequently scaled to remove cell-to-cell variation by regressing out potential technical biases. Unsupervised clustering was then performed using a graph-based algorithm implemented in the Seurat (v5.3.0) (12) package. A resolution parameter of 0.6 was applied to define distinct cell subpopulations. We utilized the genesortR (v0.4.3) (13) package to identify the top10 marker genes for each cluster based on their transcriptional profiling. Hierarchical clustering was conducted to classify the combined population of oligodendrocytes and microglia. The gene expression matrix was *Z*-score transformed (scaled) across rows to emphasize relative expression differences among clusters. The final heatmaps were generated using the pheatmap package v1.0.13 (https://CRAN.R-project.org/package=pheatmap), with hierarchical clustering performed on both rows and columns using Euclidean distance as the similarity metric and the complete linkage method.

### Differential expression analysis and functional enrichment analysis

Differential expression analysis across all microglial clusters was first performed using Seurat. For a robust comparison between the nDPM and mDPM subtypes, we employed DESeq2 (v1.46.0) (14) and limma (v3.62.2) (15). Genes meeting the criteria of *P* < 0.05 and |log2FC| > 1 were defined as significantly DEGs. The clusterProfiler (v4.14.6) (16) was utilized to conduct Gene Ontology enrichment analysis to elucidate the biological significance of these DEGs. To improve the interpretability of the results, we applied GOSemSim package (v2.32.0) (17) with a similarity cutoff of 0.8 to reduce term redundancy. Only significantly enriched terms with *P* < 0.05 were retained for further interpretation.

### WGCNA network construction and module identification

The co-expression network was constructed using the hdWGCNA (v0.4.8) (18). Quality control was first performed on the preprocessed microglia dataset, retaining genes detected in at least 5% of cells. Metacells were constructed from groups of cells sharing the same cell type and sample origin, followed by normalization of the aggregated expression matrix. After systematically evaluating different soft thresholds, a power value of 7 was selected for constructing a signed network. This power value was used to build an adjacency matrix, which was subsequently transformed into a topological overlap matrix (TOM). Co-expression modules were identified through hierarchical clustering with the grey module reserved for unassigned genes. Module eigengenes were calculated, and hub genes within each module were determined based on module connectivity (kME). Finally, module activities were assessed in the single-cell dataset using UCell scoring, based on the top 25 genes per module.

### SCENIC network construction and regulon identification

SCENIC (v1.3.1) (19) analysis was performed to reconstruct gene regulatory networks in microglia subpopulations following the established workflow. We initialized the analysis using the mm9 motif annotation database (v9) and cisTarget databases spanning 500 bp upstream and 10 kb centered on transcription start sites. Gene filtering was applied to retain genes detected in at least 1% of cells and with total counts exceeding 3% of the total cell number, yielding 8,705 genes for subsequent analysis. The log-transformed expression matrix was analyzed with GENIE3 to infer co-expression networks. Resulting TF modules were refined into regulons using RcisTarget motif enrichment with the top5perTarget method, identifying a total of 51 regulons (each containing ≥10 target genes). Regulon activity was quantified using AUCell. Regulatory intensity was visualized via Z-score normalized average AUC scores per subtype. Regulatory prevalence was assessed by binarizing AUC scores based on SCENIC-defined thresholds; only regulons active in ≥ 20% of cells in at least one subtype were visualized to evaluate the robustness of regulatory states.

### Trajectory-based mapping of microglial state to oligodendrocyte lineage

We performed pseudotemporal ordering analysis using Monocle2 (v2.34.0) (20) by integrating mDPM cells with oligodendrocyte-lineage cells. After standard normalization and the removal of lowly expressed genes (detected in < 10 cells), we selected ordering genes based on the dispersion and mean expression levels as determined by Seurat’s HVG analysis. Dimensionality reduction was performed using DDRTree and cells were ordered along the trajectory with OPC designated as the starting root.

### Cell-cell communication analysis with CellPhoneDB

Cell-cell communication was inferred using CellPhoneDB (v4.1.0) with its latest database (v5.0.0) (21). The input consisted of a normalized count matrix and corresponding metadata annotating the cell types. The statistical analysis method was executed with 1000 iterations, a p-value threshold of 0.05, and an expression threshold of 0.1 (requiring a gene to be expressed in at least 10% of cells within a cell type). The resulting interaction files, including significant means and p-values, were used for downstream visualization. Interactions were visualized using the ktplots package (v2.4.2, https://github.com/zktuong/ktplots) to generate heatmaps depicting the number of significant interactions between cell types and dot plots showing specific ligand-receptor pairs.

### Cross-dataset celltype label transferring

We collected 12 scRNAseq datasets derived from both developmental and adult hippocampus of humans and mice from both healthy and disease states (9, 22–32). For each dataset, we re-analyzed the expression matrix provided in the studies, transformed the human gene symbol or entrez ID into mouse gene symbol, and performed normalization and scaling to generate Seurat objects. Additionally, we preserved the original cell type labels each in the metadata of the Seurat object if it was provided. We kept the common genes between our data and public data, trained the SingleR model for our data by SingleR (v2.8.0) (33), and predicted the cell type of each cell of the public data by using our data as a reference.

We collected a spatial transcriptome from an adult mouse brain produced by MERFISH technology (34). Among all 59 sections, we selected the section 38 which covered most hippocampus regions. Using our data as a reference, we used CARD (v1.1) (35) to perform reference-based cell type deconvolution for the selected spatial RNAseq data.

### Feature selection and classifier construction

Data preprocessing and feature selection were performed starting with the top 100 DEGs from DESeq2 based on |log2FC|, comprising the 50 most significantly upregulated and 50 downregulated genes. Genes with an expression rate of less than 10% across all cells were filtered out. The processed dataset was partitioned into a training set (70%) and a testing set (30%) via the caret (v7.0.1) (https://github.com/topepo/caret/). For the initial feature shrinkage, LASSO regression was executed using the glmnet (v4.1.9) (36). A 10-fold cross-validation was implemented and the family set to “binomial”. The optimal penalty parameter lambda was determined by the minimum deviance criteria, and the genes with non-zero coefficients at this threshold were extracted as the characteristic feature set.

The DEGs were subsequently utilized to construct a Random Forest classifier using the randomForest package (v4.7.1.2). The model was configured with 500 trees, and the mtry parameter was optimized through a 5-fold cross-validation grid search to ensure robust performance. Variable importance was quantified using the Mean Decrease in Accuracy, identifying the key molecular features contributing to subtype discrimination. The model’s classification efficacy was rigorously evaluated on an independent testing set. Performance was visualized through Receiver Operating Characteristic (ROC) curves, with the Area Under the Curve (AUC) calculated via the pROC package (v1.18.5, https://xrobin.github.io/pROC/), and further validated by a confusion matrix to assess predictive accuracy.

### PPI Network Analysis

A protein-protein interaction (PPI) network was constructed by importing 39 overlapping genes between SCZ and AD risk genes and mDPM DEGs into the STRING database (v12.0) (https://cn.string-db.org/). The full STRING network was used with medium confidence (0.400), incorporating evidence exclusively from experiments, databases, and co-expression. The network was visualized in Cytoscape (v3.10.4) (37), where hub nodes were identified via the MCODE plugin based on Degree centrality (38). Node size and color gradients were mapped to degree scores to highlight core functional clusters.

### Reads analysis

We extracted reads of DPM cells, classifical microglial cells and cells of OL lineage (including OPC, OPC_variant, COP, oligo_classical and oligo_variant) from the bam files. We then used velocyto^59^ to calculate the unspliced reads count (#u) and spliced reads count (#s) with “run-dropest” mode and “Stricter10X” logic. The genome annotation file used in velocyto analysis is of GRCm38.97 version. Each sample was processed separately, and the result loom files were merged by combine function of loompy (v3.0.8) package (https://github.com/linnarsson-lab/loompy). The unspliced reads ratio of gene was defined as the ratio of #u to total reads count. The genome browser visualization of St18 was performed by IGV (v2.19.7) (39).

### Generative AI Usage

The authors utilized Google Gemini and Deepseek as assistive tools to enhance the readability of the text and to optimize the presentation of the scientific figures. All AI-generated suggestions were critically evaluated and integrated into the manuscript by the authors.

## Results

### Two distinct DPM subtypes with diverse biological functions were identified

Based on our previously published single-cell transcriptomic data of the hippocampus from adult mice, we extracted the microglial population from wild-type mice for deeper analysis(11). To further explore microglial heterogeneity, we performed subclustering on microglial cells and obtained five cell clusters. Cluster 0 and 1 represented classical microglia, while Cluster 4 corresponded to a macrophage population. Clusters 2 and 3 co-expressed established microglial and oligodendrocytic markers, designated as DPM population (Figure 1A-B, Additional file 1: Figure S1A). In contrast to the highly similar transcriptomes of Clusters 0 and 1, Clusters 2 and 3 displayed markedly divergent transcriptional profiles, underscoring that the DPM was segregated into two discrete cell subtypes (Figure 1C; Additional file 2: Table S1).

**Figure 1.**
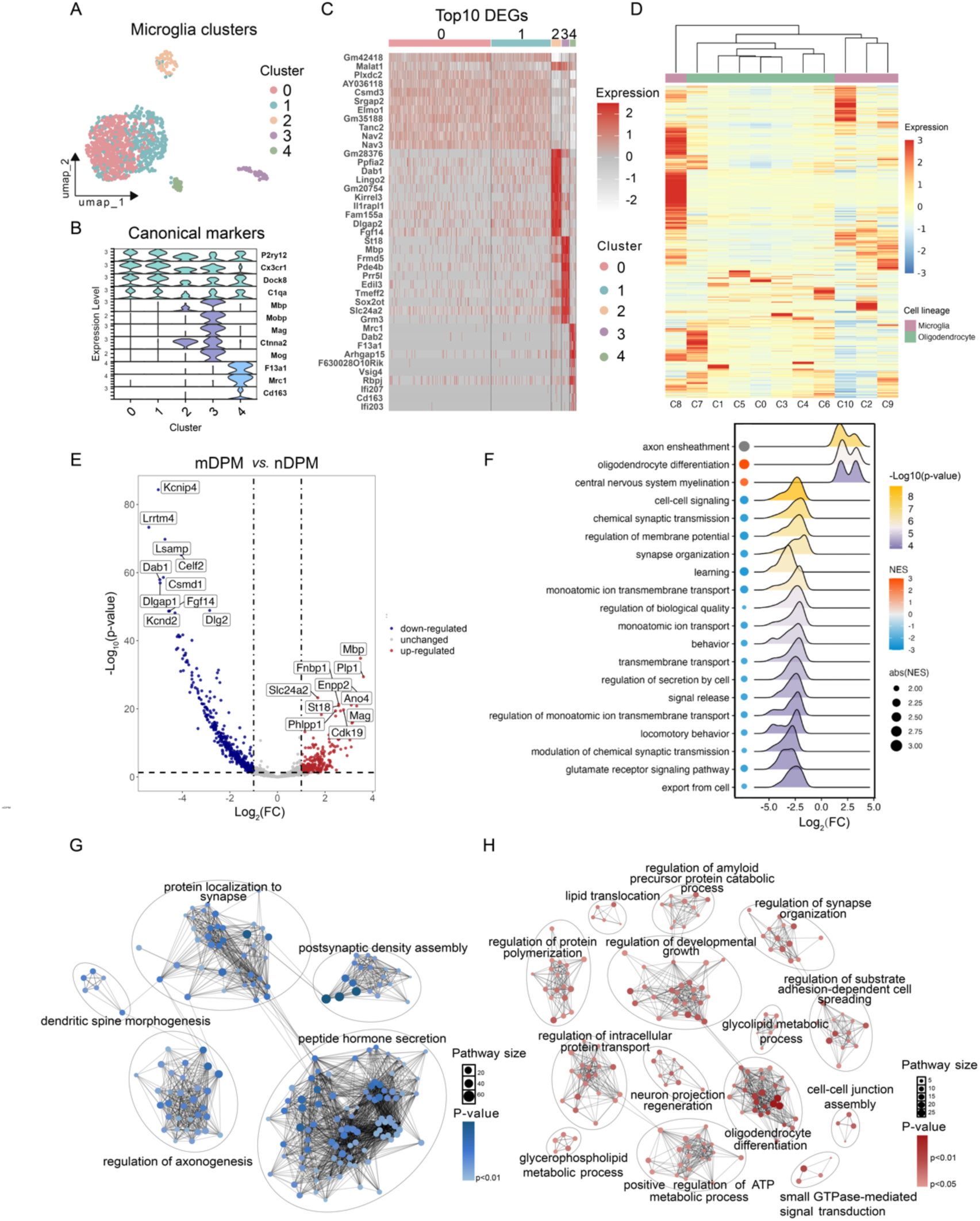
Identification of DPM subtypes. **(A)** UMAP visualization of classical microglia and DPM clusters. **(B)** Violin plots displaying the expression levels of canonical markers across clusters. **(C)** Heatmap showing the top 10 DEGs for each cluster identified by genesorteR. **(D)** Hierarchical clustering and heatmap of scaled gene expression across all identified clusters. **(E)** Volcano plot comparing mDPM (Cluster 3) and nDPM (Cluster 2) by DESeq2. Log2 fold change > 1 and *P <* 0.05. **(F)** GSEA ridge plot. Points and density curves show the distribution of gene fold changes within specific GO terms. **G–H:** Enrichment-based functional networks for nDPM (G) and mDPM (H). Nodes represent GO terms and edges denote shared gene membership between terms.

Given the expression of oligodendrocytic markers in DPMs, we sought to clarify their relationship with canonical oligodendrocyte lineages. Combining oligodendrocytes, classical microglia, and DPMs for unsupervised clustering analysis generated 11 subclusters. Hierarchical clustering based on average gene expression revealed that the two DPM clusters—C8 (corresponding to Cluster 2 in Figure 1C) and C9 (corresponding to Cluster 3 in Figure 1C)—were separated in the hierarchical tree (Figure 1D, Additional file 1: Figure S1B). C9 exhibited a gene expression profile more closely aligned with classical microglia, whereas C8 occupied an intermediate position, distinct from both classical oligodendrocytes and microglia. This divergence indicateed potentially different cellular origins for the two DPM subtypes.

To demarcate and define DPM subtypes, we performed gene ontology (GO) enrichment analysis. Cluster 2 was significantly enriched in terms related to “dendrite development”, “cognition”, and “regulation of synapse structure or activity” implicating a role in neuronal processes; we thus designated this subtype as neuron-associated DPM (nDPM). In contrast, Cluster 3 was enriched for biological functions such as “glial cell differentiation”, “axon ensheathment”, “ensheathment of neurons”, and “myelination” supporting its classification as myelin-associated DPM (mDPM) (Additional file 1: Figure S1C). To delineate the molecular distinctions between mDPM and nDPM, we performed differential expression analyses using limma and DESeq2, both analysis strategies consistently identified robust transcriptional differences between the two subtypes (Additional file 2: Table S2–S3). The mDPM specifically upregulated genes encoding key myelin structural components (e.g. *Mbp*, *Plp1*, *Mag*) (40) and the oligodendrogenic transcription factor *St18* (41). Contrastively, nDPM was enriched for genes involved in neuronal development and synaptic regulation, including *Dab1* (essential for neuronal migration during development) (42), *Csmd1* (Complement inhibitor in developmental synaptic pruning) (43), and *Ppfia2* and *Dlgap1* (regulators of synaptic plasticity) (44, 45) (Figure 1E, Additional file 2: Figure S1D). Together, both differentially expressed genes (DEGs) and GO analyses established a foundational definition of the two functionally distinct DPM subtypes.

To further explore the biological processes and the interconnected function of these subtypes, we performed Gene Set Enrichment Analysis (GSEA). Consistent with the individual gene markers, the GSEA results reinforced that mDPM is driven by coordinated programs of axon ensheathment, oligodendrocyte differentiation, and CNS myelination. For nDPM, the analysis revealed the enrichment in processes governing neuronal communication, such as chemical synaptic transmission, membrane potential regulation, and glutamate receptor signaling (Figure 1F). To visualize the high-order organization of these functional modules, we constructed enrichment-based networks (Figure 1G–1H). These networks revealed that the nDPM-associated terms form tightly coupled clusters centered on synaptic assembly and regulation of axonogenesis, while the mDPM is characterized by integrated modules of oligodendrocyte differentiation and developmental growth.

### Bioinformatic and immunofluorescent approaches validated the spatial distribution pattern of DPM subtypes

Considering the prominent distinction of cellular function of DPM subtypes, we further explored the spatial distribution pattern of them. We mapped our scRNA-seq-defined microglial clusters onto whole-brain spatial transcriptomic data from adult mice(34). The predicted cell type across the brain regions elucidated high specificity of the spatial distribution pattern of two DPM subtypes. Specifically, nDPM was significantly enriched in the hippocampal dentate gyrus (DG)—a key niche for adult neurogenesis (Figure 2A, left panel). This spatial localization was consistent with its functional indication transcriptional signature related to neuronal development and synaptic regulation, suggesting a potential role in modulating the survival, differentiation, or integration of newborn neurons. By contrast with nDPM, mDMP exhibited a sparse distribution pattern across the entire brain (Figure 2A, right panel), suggesting that they were not restricted to any specific anatomical compartment. This pan-brain presence, combined with a molecular profile enriched for ‘axon ensheathment’ and ‘myelination’, indicated that mDMP-mediated regulatory functions were fundamental and widespread, potentially serving to maintain axonal homeostasis.

**Figure 2.**
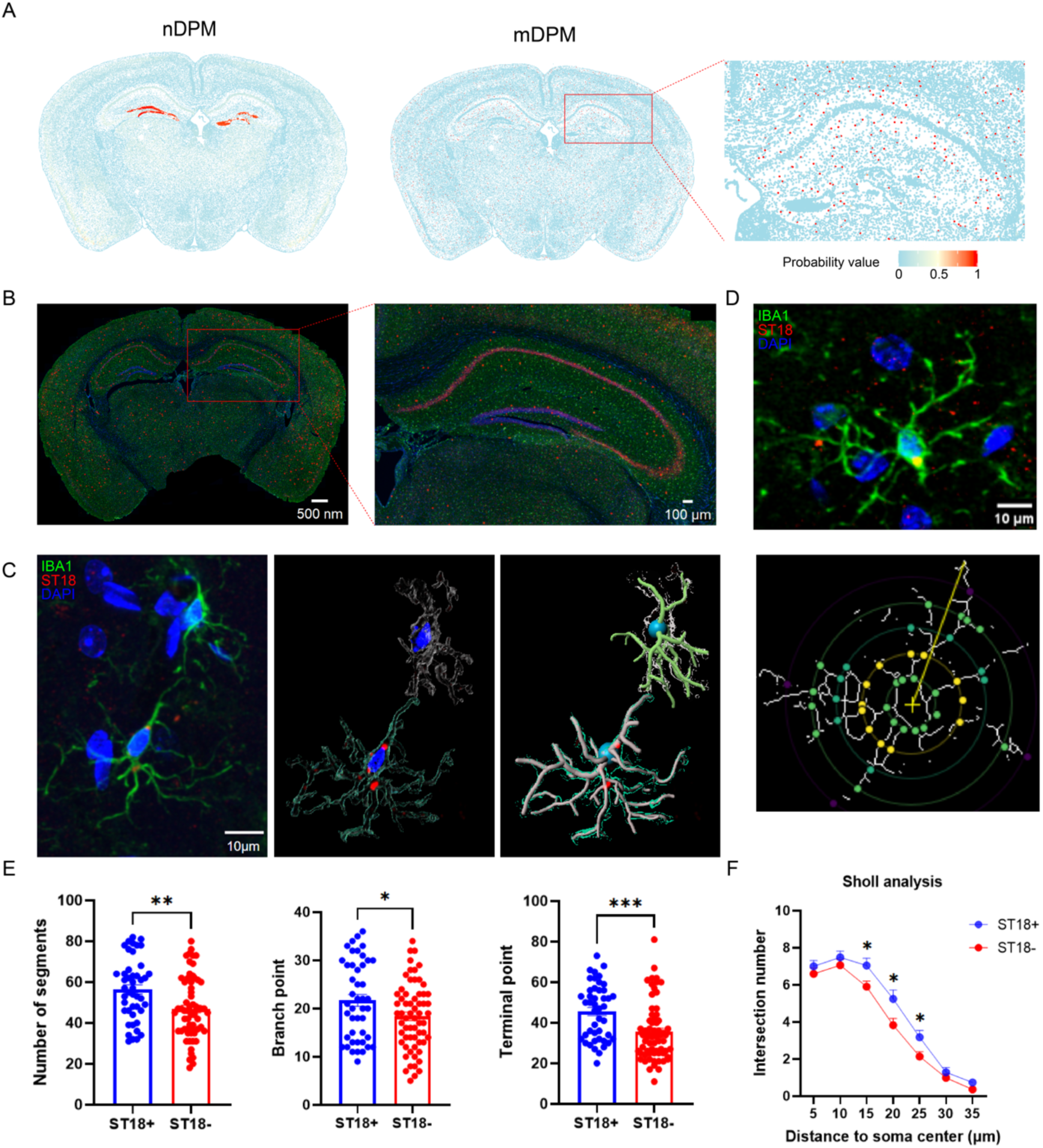
Spatial distribution and morphological characterization of DPM subtypes. **(A)** Spatial mapping of nDPM (left) and mDPM (middle) subtypes onto adult mouse brain sections using spatial transcriptomics data. A magnified view of the hippocampus (right) shows the enrichment of mDMP in this region. **(B)** Representative whole-brain immunofluorescence image (left) and magnified hippocampal section (right) showing ST18 expression (red) in Iba1-labeled microglia (green). **(C)** Representative 3D reconstruction of ST18+ (mDPM) and ST18-microglia. Left: Original IF image; Middle Surface rendering; Right: Skeletonized filament model for morphometric analysis. **(D)** Representative image of immunofluorescence (top) and 2D Sholl analysis overlay (bottom) used for morphological quantification. **(E)** Quantitative comparison of morphological parameters derived from 2D reconstruction between ST18+ (n = 45) and ST18– (n = 64) microglia. Data are presented as mean ± SEM; unpaired t-test, *P < 0.05, **P < 0.01, ***P < 0.001. **(F)** 2D Sholl analysis curves showing the number of intersections at increasing distances from the soma center for ST18+ and ST18-microglia.

To confirm the biological function of the DPM subtypes robustly, we involved two independent public datasets, and compared our findings with the cell types with similar cellular attributes. First, a microglial population from a human hippocampal single-nucleus dataset (9) with functional enrichment of “myelination” and “glial cell differentiation”, showed high expression of *PKP4* and *MYO1D*, which were the marker genes of mDPM (Additional file 1: Figure S2A). Second, we extracted the transcriptional signature of Myelin-transcript-enriched-microglia (MyTE) from a mouse model (46); scored the single cells in our dataset with the marker geneset of MyTE, and found that the mDPM subtype exhibited significantly higher scores compared to all other microglial subtypes (Additional file 1: Figure S2B). These results underscored the robustness of DPM across species and experimental systems, suggesting that the DPM cells are stable cell type rather than a dataset-specific artifact.

Apart from the cross-datasets and cross-species evidence from informatic approach, we sought to determined the representative markers of DPM subtypes and performed experimental validation. Firstly, to identify the most discriminative molecular features distinguishing two DPM subtypes, we employed LASSO regression and Random Forest (RF) algorithms to analyze the top 50 upregulated and top 50 downregulated DEGs between nDPM and mDPM from the results of DESeq2 analysis (Additional file 2: Table S4). The LASSO and RF models selected 17 and 30 features respectively, based on which, we defined a consensus feature set by using 12 overlapped genes of the results from two machine learning models (e.g., *Dlgap1*, *Dab1, Csmd1*) (Additional file 1: Figure S2C–S2D). Both models exhibited high discriminative capacity, consistent with the transcriptomic results observed between the two clusters. Notably, these top genes were predominantly nDPM-upregulated genes involved in synaptic function and neuroelectrical signaling. This bias toward nDPM markers likely stemed from the fact that their fold-change magnitudes were substantially greater than those of myelin-related genes in mDPM, providing maximal separability for the algorithms.

The above part enabled us refine the DE gene set into a core diagnostic signature. Next, we selected the top significant feature of nDPM and mDPM from the 12 consensus feature set for in-situ validation. Considering the availability of validated commercial antibodies, we selected DAB1 (a signature gene for nDPM linked to neuronal migration) (42) and ST18 (a specific marker for mDPM essential for myelination) (41) as representative markers to perform immunofluorescence in brain tissues of adult male mice. Consistent with the spatial cell mapping pattern, St18+ microglia (representing mDPM) were broadly distributed across the whole brain (Figure 2B). To further determine the structural features of ST18+ microglia, we performed high-resolution three-dimensional (3D) reconstruction and Sholl analysis to compare the morphology of ST18+ microglia with that of ST18-microglia (Figure 2C). The 3D analysis revealed that mDPM possessed a significantly more elaborated architecture than classical microglia. Specifically, ST18+ microglia exhibited higher ramification, with a significant increase in the number of segments (*P* < 0.01), branching points (*P* < 0.05) and terminal points (*P* < 0.001) (Additional file 1: Figure S2E). This structural complexity was further corroborated by the two-dimensional (2D) Sholl analysis, which consistently demonstrated an increase in branching density in the mDPM subtype (Figure 2D–2F). In contrast, DAB1+ microglia (nDPM) exhibited a morphology indistinguishable from DAB1-counterparts across all metrics and Sholl analysis, suggesting that, unlike mDPMs, nDPMs did not undergo specialized structural alteration under homeostatic conditions (Additional file 1: Figure S2F).

In summary, the in-situ validations confirmed that ST18 and DAB1 were reliable molecular identifiers for mDPM and nDPM, respectively, and highlighted the distinct morphological and spatial attributes that defined these two functional states.

### Functional analyses emphasized molecular-level signatures of DPM subtypes

Gene-level analyses have suggested the functional specificity of mDPM and nDPM. To systematically characterize the molecular programs governing the functional specialization of DPM subtypes, we performed a gene module-level analysis using weighted gene co-expression network analysis (WGCNA). According to the pair-wised correlation of their expression level, single genes were grouped into several distinct co-expression modules, each showing preferential association with different DPM subtypes (Additional file 1: Figure S3A). Module1 exhibited a strong and specific association with nDPM. GO enrichment analysis revealed that genes within this module were significantly involved in nervous system development and cell-cell signaling (Figure 3A–3B). Notably, several hub genes in Module 1 were established regulators of neuronal structure and function, including *Rbfox3* (a mature neuron marker, also known as NeuN), *Dab1*, *Dlgap1* (a postsynaptic scaffolding protein) (47), alongside neurotransmitter receptors *Gabrb2* and *Grik2*. The coordinated expression of these functionally related genes indicated that nDPM possessed a coherent transcriptional program specialized for involving in neuronal developmental and synaptic processes.

**Figure 3.**
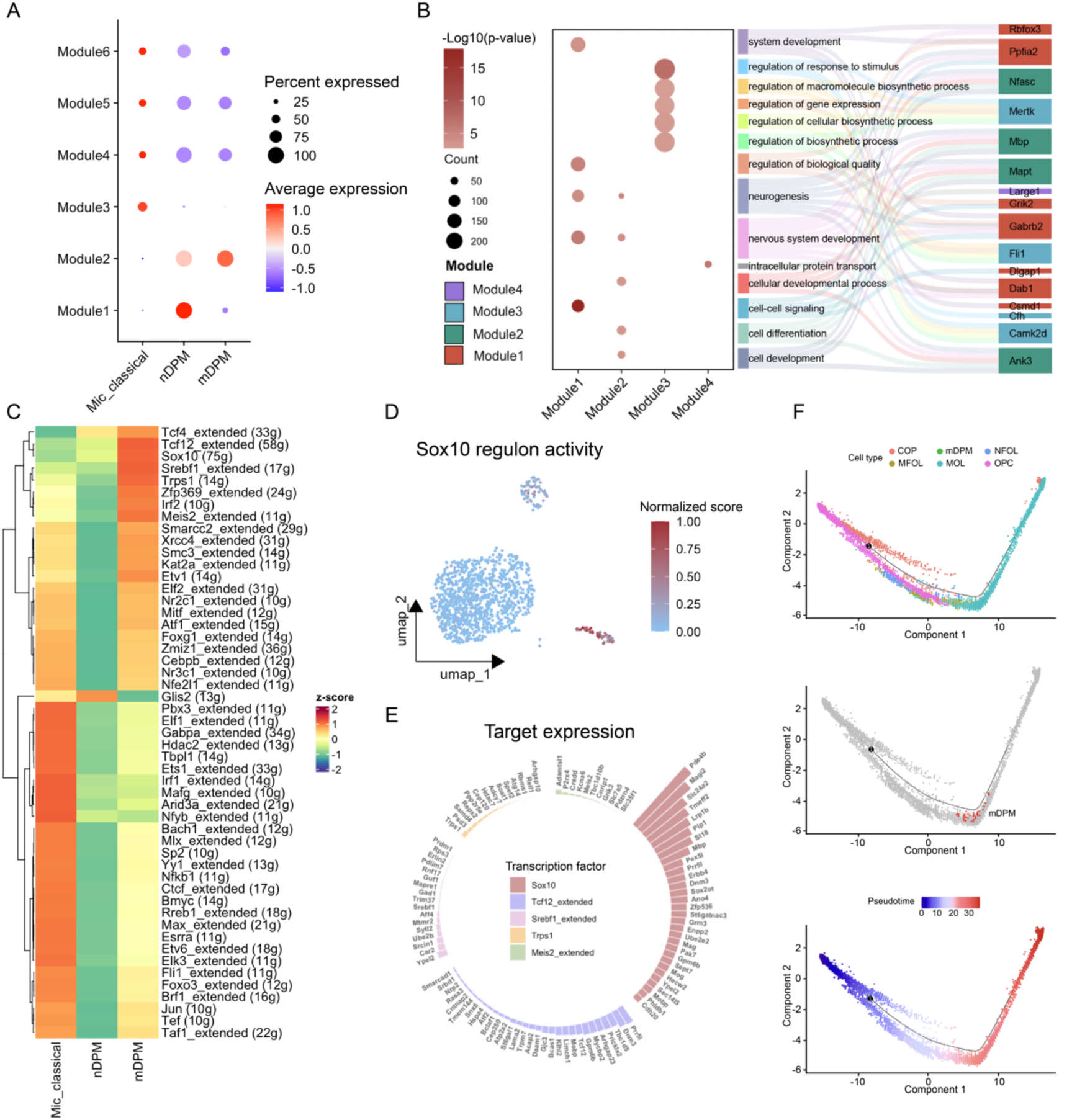
Co-expression network and transcription factor regulatory network of DPM subtypes. **(A)** Average expression levels of six co-expression modules across classical microglia, nDPM, and mDPM subtypes. **(B)** Functional enrichment and hub genes of modules. **(C)** Heatmap showing the average regulon activity scores (derived from AUCell) for each microglial subtype. Data are presented as row-scaled Z-scores. **(D)** UMAP visualization showing the normalized enrichment score of the *Sox10* regulon. **(E)** Circular visualization of mDPM-specific regulatory networks. Target genes controlled by the top five regulons enriched in mDPM. **(F)** UMAP plots showing the integration of DPM subtypes with the oligodendrocyte lineage (top), the distribution of cell types (middle), and the inferred pseudotime (bottom).

To further explore this potential, we examined the intercellular communication using CellPhoneDB (21), focusing specifically on ligand-receptor pairs between nDPM and both excitatory and inhibitory neurons. We performed separate functional enrichment analyses for ligands and receptors exclusively expressed by nDPMs within these identified pairs. When acting as a signal sender (analysis based on nDPM-expressed ligands), processes associated with neuronal growth and synaptic maturation were enriched in nDPM, including “cell growth”, “developmental growth involved in morphogenesis”, “synapse assembly”, and “axonogenesis”. When functioning as a signal receiver (analysis based on nDPM-expressed receptors), nDPM exhibited enrichment in “neuron projection guidance” and “G protein-coupled receptor signaling” (Additional file 1: Figure S3B–S3C). These spatially resolved signaling profiles reinforced the notion that nDPM was molecularly equipped to both modulate the neuronal structural maturation and respond to cues from the local neuronal environment.

We then investigated the transcription factor (TF)-target networks by SCENIC(19).The analysis identified regulons, which were functional modules comprising a TF and its co-expressed targets supported by motif enrichment, to map the upstream regulation. We found that *Glis2* regulon was significantly enriched in nDPM (Figure 3C). Consistent with the role of *Glis2* in neuronal differentiation(48), we found that its downstream target, *Zhx2*—a known maintainer of neural progenitor states (49)—was also specifically upregulated in nDPM (Additional file 1: Figure S3D). Together, these findings suggested that a *Glis2* centered transcriptional network may contribute to the molecular features of nDPM associated with neuronal development related processes.

In parallel, mDPM showed a preferential association with Module2 (Figure 3A), which was enriched for cell differentiation and myelination-related processes. Several hub genes within this module were critically linked to myelination and axon-glial interactions. These included *Nfasc*, which mediates axon-glial adhesion at the paranodal region for node of Ranvier formation (50); *Mapt* (encoding Tau), a protein known to promote myelination via FYN kinase signaling (51); and *Ank3* (encoding Ankyrin G), a scaffold protein essential for clustering sodium channels at axonal segments (He et al., 2022). The expression of these genes, alongside the core myelin component *Mbp* (52), suggested that mDPM may actively engage in the maintenance and organization of specialized axonal domains (Figure 3B). Notably, despite acquiring this myelin-associated profile, mDPM cells consistently retained core microglial markers (e.g., *Cx3cr1*, *C1qa*, and *P2ry12*), maintaining their distinct identity from the macroglial lineage (Figure 1B).

Transcription factor regulon analysis further identified a pronounced enrichment of the *Sox10* regulon within the mDPM (Figure 3C). To ensure the regulatory programs were representative of the entire subtype rather than driven by outliers in a few cells, we examined the binarized regulon activity, which confirmed that the *Sox10* regulon was active in a high proportion of mDPM cells, supporting the robustness of this lineage-specific transcriptional state (Additional file 1: Figure S3E). *Sox10* is a master regulator of oligodendrocyte differentiation and its regulon encompassed canonical myelin genes such as *Plp1*, *Mbp*, and *Mag*. Gene set scoring and GO enrichment confirmed that the *Sox10* regulon and its target genes exhibited significantly higher activity in mDPM and were primarily associated with axon ensheatment and oligodendrocyte development (Figure 3D–3E, Additional file 1: Figure S3F; Additional file 2: Table S5). These findings indicated that mDPM engaged a *Sox10* centered transcriptional program that closely resembled the signature of the myelination and the structural maintenance of the axon-glial interface.

In order to position mDPM relative to the oligodendrocyte (OL) differentiation trajectory, we integrated our dataset with a published hippocampal atlas (53) comprising a full continuum of OL maturation stages. And then we performed a joint pseudotime analysis with mDPM, OPCs, COPs, NFOLs, MFOLs and MOLs. The analysis revealed that mDPM mapped to a transcriptional standing corresponding to myelinating precursor oligodendrocytes (MFOLs) (Figure 3F), which was consistent with their active transcriptional state in myelination related functions. This convergence along the oligodendrocyte developmental trajectory, coupled with the retention of microglial identity, suggested that mDPM occupied a unique functional state that balanced microglial identity with an active transcriptional program typically reserved for myelination.

### Alteration of DPM-like cells underlied neuropsychiatric disorders

To define the pathological relevance of DPM subtypes, we intersected their differentially expressed gene signatures with risk gene lists from the Phenopedia database v10.0(54). This analysis revealed that nDPM-specific genes were significantly enriched for schizophrenia (SCZ), major depressive disorder (MDD), and autism spectrum disorder (ASD) (Figure 4A). In contrast, mDPM genes showed specific enrichment for SCZ and Alzheimer’s disease (AD) (Figure 4A), both diseases being characterized by white matter abnormalities and progressive myelin loss(55–57), suggesting a potential association between mDPM and pathologies involving myelination and axonal integrity. To decipher the molecular pathways underlying the genetic associations, we performed enrichment analysis on the intersecting gene sets. For the genes shared between mDPM and AD, pathways including “amyloid-beta binding”, “APP metabolic process”, and “positive regulation of myelination” were significantly enriched (Figure 4B). Similarly, analysis of the mDPM-SCZ shared genes revealed association with “structural constituent of myelin sheath”, “glutamate receptor activity”, and “postsynapse organization”, reflecting the co-dysregulation of myelination and synaptic signaling in SCZ (Figure 4C).

**Figure 4.**
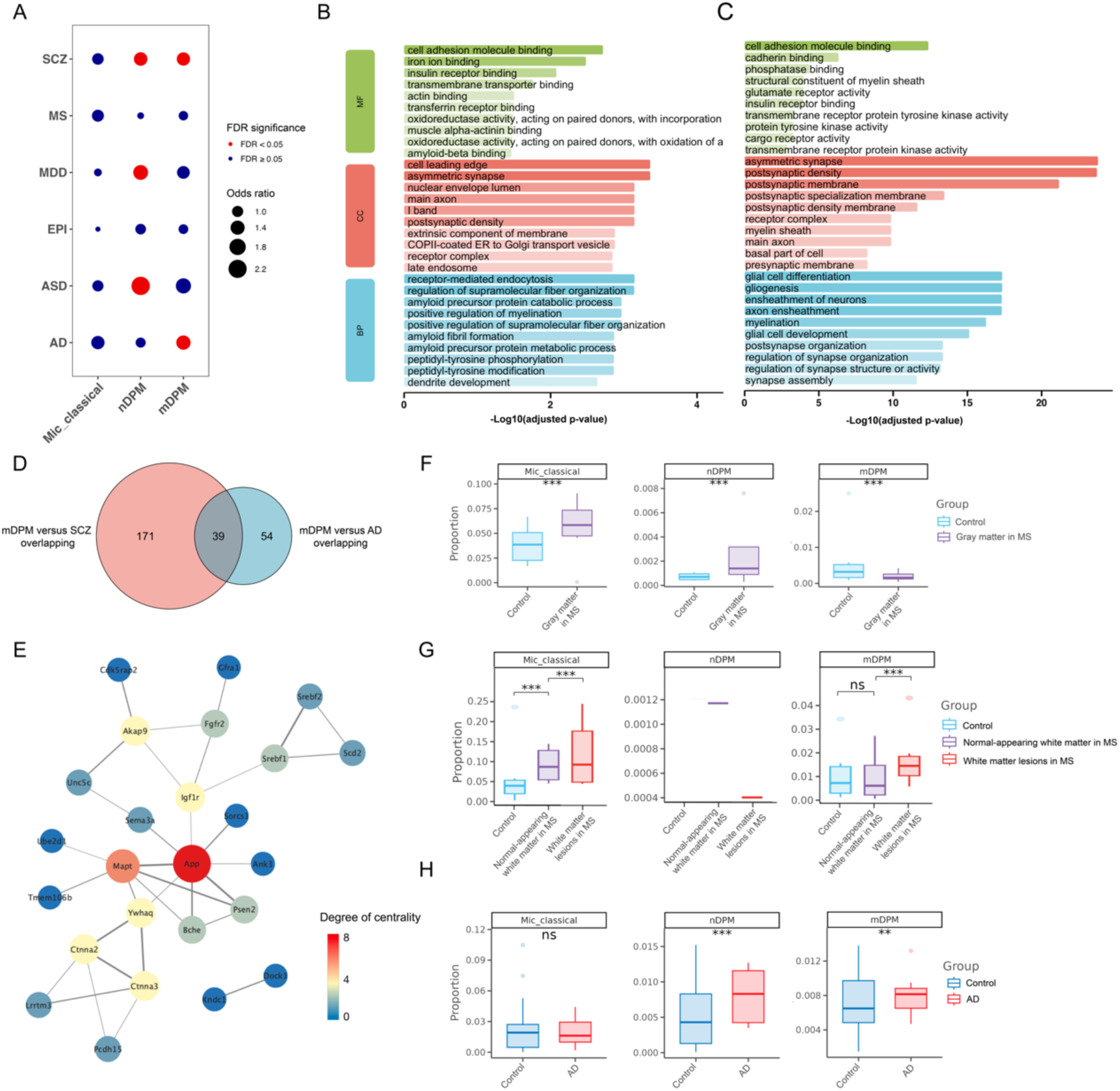
Characterization of disease features and celluar dynamics in DPM Subtypes. **(A)** Disease-risk gene enrichment analysis of DPM subtypes. Statistical significance was determined by Fisher’s exact test using human-mouse orthologous genes as background. Disease-risk genes were obtained from the Phenopedia database. **(B–C)** Bar plots showing enriched pathways for genes intersecting mDPM with AD (B) and SCZ (C). **(D)** Intersection of mDPM-enriched DEGs with risk gene sets for SCZ and AD. Nodes are sized and colored according to their degree centrality, with larger red nodes representing high-degree hub genes and smaller blue nodes representing lower-degree genes. **(E)** Interaction network of the 39 genes shared among mDPM, AD, and SCZ. **(F–H)** DPM proportions in human diseases. Proportions of DPM subtypes in MS gray matter (F), MS active lesions (G), and AD hippocampus (H). Statistical significance determined by two-sample proportions test followed by Benjamini-Hochberg (FDR) correction.

Further investigation into the genetic overlap between mDPM, AD, and SCZ identified 39 common genes (Figure 4D). A protein-protein interaction (PPI) network of these shared genes highlighted APP and MAPT (tau) as central hubs (Figure 4E). Notably, these hubs exhibited dense connections with key growth factor receptors such as IGF1R and FGFR2. Given the established roles of IGF and FGF signaling in microglial-mediated remyelination and oligodendrocyte support (58), the central position of these molecules suggested that the genetic risk associated with mDPM may converge on growth factor-mediated pathways essential for axonal ensheatment and myelin maintenance.

Parallel enrichment analysis of the gene sets intersecting nDPM with each of its associated disorders (SCZ, MDD, and ASD) showed that, despite their distinct clinical contexts, all three gene sets converged on core pathways related to synaptic function and processes regulating neuronal signaling (Additional file 1: Figure S4A–S4C). A Venn diagram analysis identified a consensus set of 122 genes that were both differentially expressed in nDPM and underlied the genetic risk for SCZ and MDD (Additional file 1: Figure S4D). This genetic convergence strongly reinforced nDPM’s role as a specialized regulator of neuronal communication and suggested that this 122-gene core may represent a shared molecular substrate for nDPM-mediated synaptic dysfunction across multiple psychiatric conditions.

We next examined whether these genetic associations corresponded to alterations in DPM abundance within diseased human tissue. Multiple Sclerosis (MS), a disease defined by targeted inflammatory attacks on myelin sheaths and axonal degeneration, provides a highly relevant pathological environment to assess the response of mDPM (59). Analysis of single-cell datasets from patients with multiple sclerosis (MS) (25) revealed region-specific alterations. In the gray matter, the proportion of predicted-mDPM was markedly reduced, whereas predicted-classical microglia and predicted-nDPM were increased (Figure 4F). In the white matter compartment, classical microglia were significantly elevated compared to controls, while predicted-mDPM levels remained comparable. Notably, direct comparison within MS tissue samples demonstrated that mDPM was selectively enriched in active demyelinating lesions relative to the adjacent plaque white matter, indicating its targeted recruitment to sites of active pathology (Figure 4G). Furthermore, in the hippocampus of AD patients (9), both predicted-nDPM and predicted-mDPM populations were significantly expanded relative to controls, while classical microglia remained unchanged (Figure 4H). The distinct pattern of predicted-mDPM depletion in MS gray matter, in contrast to its enrichment in the AD hippocampus and within active MS lesions, underscored their plastic and potentially active roles in responding to pathological state.

To understand the significance of these disease-specific alterations, we contextualized them within the normal physiological framework of DPM biology by assessing their dynamics throughout development. Both subtypes represent sustained cellular states within the microglial repertoire but display stage-specific patterns. mDPM peaked during early embryogenesis (E11), while nDPM reached its maximum proportion during the late embryonic period (E16.25) (Additional file 1: Figure S4E). In the postnatal hippocampus, nDPM abundance coincided with phases of active synaptic refinement (60), while mDPM increased during adulthood and aging, aligning with periods of ongoing myelin plasticity (Additional file 1: Figure S4F). Together, these physiological benchmarks indicated that the reconfiguration of DPM proportions observed in MS and AD reflected a specific, disease-driven reorganization. The enrichment of mDPMs within active MS lesions and the AD hippocampus points to a targeted mobilization through recruitment from unaffected areas or local expansion.

## Discussion

In this study, we comprehensively described and explored the biological function and morphological phenotype of a specific subtype of microglial cells, DPMs. The presence of this cell type has been reported in our previous study as well as other research(8, 9, 46). However, in the present study, we sought to provide highly reliable evidence that these cells constitute a genuine, stable cell type. Apart from the informatic and experimental validation (see Figure 1**–**3), we further analyzed the sequencing reads of DPM marker genes to rule out other potential biological confounding factors, such as the phagocytic activity of microglial cells, which was mentioned in Schirmer L et al.’s study (61). The authors suggested that microglial enrichment for oligodendrocyte-specific markers *Plp1*, *Mbp*, and *St18* could reflect phagocytosis of myelin-derived RNA.

To rule out the possibility of a contribution from microglial cell phagocytosis, we extracted the reads of DPM cells from sorted bam files, implemented velocyto analysis to determine the reads as unspliced, spliced and ambigious by Stricter10X logic. The Stricter10X logic has more strict criteria that only non-singletons supported by validated introns are counted as unspliced. The unspliced ratio of gene was calculate as the ratio of unspliced reads count to the total reads count. We found that mDPM showed higher unspliced ratio in *St18* (mean value = 0.98) compared to that of oligodendrocytes (mean value is 0.96). By contrast, other markers of oligodendrocytes such as *Olig2*, *Cnp* and *Apc* had lower ratio in mDPM, while for gene like *Slca17a7* that is not marker of neither mDPM nor oligodendrocytes we barely observed unspliced reads (Additional file 2: Table S6). Take *Olig2* as positive control and *Slca17a7* as negative control, we conclude that mDPM expressed *St18* natively rather than phagocytose the *St18* products of oligodentrocytes. Additionally, we visualize the distribution across *St18* genic region of reads of mDPM by IGV tool, especially in intronic regions (Additional file 1: Figure S5).

The discovery redefined the mechanism by which microglia acquire oligodendrocyte-associated transcripts. While previous studies frequently attributed such phenomena to passive phagocytosis by positioning microglia solely as scavengers of myelin debris, our evidence suggested a more sophisticated biological role. This conclusion was further corroborated by geneset enrichment scoring for phagocytic pathways (Additional file 1: Figure S1E), which revealed that traditional phagocytic machineries were not activated within these specific subtypes.

In further analysis, hierarchical clustering analysis suggested potentially divergent developmental origins for these two subtypes, with mDPMs exhibiting a closer transcriptomic proximity to the classical microglial lineage. mDPMs displayed a widespread distribution across the brain parenchyma (Figure 2A–2B). Despite this scattered localization, mDPMs maintained a unique oligodendrocyte-like transcriptional program that confers a distinct functional identity. Our data demonstrated that mDPMs were not merely recycling environmental RNA but were actively executing an endogenous, lineage-specific regulatory program. This functional state appeared to be driven by a reconfigured transcriptional framework, notably involving the transcription factors *Sox10* and *St18*. Of particular interest is *St18*, which, as an inhibitory regulator, may function as a critical ‘molecular switch’ governing the state transition of mDPMs and controlling the ectopic expression of myelin-associated genes.

Given that the myelin sheath is a highly dynamic structure, its functional integrity strictly depends on continuous lipid turnover and metabolic equilibrium(62). The persistent presence of mDPMs across all life stages, their highly ramified morphology, and their specific enrichment in myelin-metabolic genes (e.g., *App*, *Lpl*) strongly suggest that mDPMs function as auxiliary glial units dedicated to supporting myelin homeostasis, rather than acting as transient immune responders.

The above deduction together with the mDPM gene signature (Figure 1C, 3B–3C) provided the inspiration about the picture of mDPM managing the complete cycle of myelin lipid turnover: Specifically, mDPMs operate as a specialized “metabolic relay” by adopting a trans-lineage transcriptional program driven by master regulons *Sox10*, *Tcf12*, and *Srebf1*. While maintaining a ramified, surveillance-ready morphology through *Frmd5*, these cells express a suite of oligodendrocyte-specific structural genes—including *Mbp*, *Plp1*, *Mag*, *St18*, and the regulatory RNA *Sox2ot* —to enable precise, “same-language” communication with the myelin sheath. Beyond simple phagocytosis, they actively manage lipid homeostasis via *Enpp2* and *Srebf1*, and potentially provide structural scaffolding at the axon-glial junction through hub genes like *Mapt* and *Ank3*. By integrating environmental sensing through *Grm3* and *Slc24a2*, anti-inflammatory control via *Pde4b*, and vesicle-mediated support through *Dnm3*, alongside auxiliary regulators like *Prr5l*, *Edil3*, and *Tmeff2*, mDPMs establish a comprehensive functional backup that secures myelin integrity and metabolic stability.

In steady-state conditions, they appeared to serve as homeostatic sentinels that monitor and sense the myelin microenvironment, actively participating in the precise regulation of lipid turnover and routine myelin maintenance. While in the context of demyelinating diseases, this homeostatic balance was disrupted. As mDPMs transitioned from steady-state sentinels to pathological participants, their pro-myelinating programs may undergo maladaptive polarization. In MS, the selective enrichment of mDPMs in active lesions likely reflected a compensatory reparative effort, an attempt to rescue the damaged myelin environment by expanding this specialized subtypes. However, whether chronic pathological stress eventually leads to the functional exhaustion of this mimicry program, thereby exacerbating demyelination, remains a critical question.

Furthermore, interaction analysis of mDPM-specific risk genes revealed a significant enrichment in genetic susceptibility for SCZ and AD. Notably, these shared risk genes converged on APP and MAPT (Tau) as central hubs within the protein-protein interaction (PPI) network, exhibiting dense functional connectivity with key growth factor receptors such as IGF1R and FGFR2. Given the established roles of IGF and FGF signaling as cornerstones for microglial-mediated remyelination and oligodendrocyte maturation (63), the centrality of APP and Tau suggested that genetic risk in these disorders may converge on growth factor-mediated pathways essential for axonal ensheatment and myelin stability. The risk genes proposed a pathological hypothesis: in the early stages of disease, abnormalities in APP or Tau may first impair the sentinel function of the mDPM subtypes. This disruption of axonal surveillance and maintenance could lead to a chronic decline in myelin integrity, ultimately impairing the connectivity of long-range neural circuits—a potential cellular foundation for the cognitive and psychiatric symptoms observed in AD and SCZ.

The significance of covering the sophisticated biological states (e.g. DPM) is further highlighted when contextualized within recent macro-scale transcriptomic landscapes (64, 65), which established comprehensive microglia taxonomy. To achieve such unprecedented scale and cross-condition consistency, large-scale atlases inherently employ highly stringent doublet-filtering pipelines and rely on classical mononuclear phagocyte boundaries to ensure lineage purity. While this methodology is highly effective for defining core immune populations, our findings suggest that strictly penalizing cross-lineage co-expression may inadvertently mask highly plastic, transitional cell states. Rare biological entities like DPMs, which endogenously co-express microglial and oligodendrocyte markers, might be algorithmically filtered out as technical doublets or merged into broad “phagocytosis” superclusters. Therefore, our study provides a vital complementary perspective to these expansive atlases. By utilizing RNA velocity to confirm *de novo* transcription and 3D morphological reconstruction, we highlight the vertical depth of microglial plasticity. Our work demonstrates that capturing these subtle, trans-lineage reprogrammings is crucial for fully understanding the diverse compensatory mechanisms of microglia in health and disease.

Several limitations in this study warrant further discussion. A key unresolved question is the developmental origin of these two DPM subsets. We currently cannot discern whether mDPMs and nDPMs arise from independent cell lineages or if they are simply different functional states governed by complex epigenetic regulation upon brain colonization. This possibility is underscored by evidence that microglia possess significant epigenetic plasticity and drives microglia-to-neuron conversion (66). However, whether the divergence between nDPMs and mDPMs is similarly rooted in specific chromatin accessibility changes or histone modifications requires further validation by epigenomics data. Furthermore, although we identified a clear redistribution of mDPMs in both MS and AD, the specific associations between these cells and disease progression have not yet been deeply explored. Compelling empirical evidence is still required to definitively establish the functional contributions of mDPMs to these pathological states. Ultimately, the hypothesized association between mDPMs and the oligodendrocyte-myelin unit awaits experimental confirmation. Future work using spatial metabolomics or lineage tracing will be vital to verify if mDPMs indeed serve as active participants in myelin lipid turnover as we propose.

## Conclusions

In conclusion, we present robust transcriptomic, spatial, and morphological evidence for the existence of Dual-Phenotype Microglia (DPM), including Myelin-associated DPM (mDPM) and Neuron-associated DPM (nDPM)—two microglial subpopulations that intrinsically executes oligodendrocyte– and neuron-associated transcriptional programs while retaining its core microglial identity. Furthermore, we establish the profound clinical relevance of these subsets, with mDPMs being selectively recruited and enriched in the active demyelinating lesions of MS and the hippocampus of AD patients, and mDPM-specific gene signatures show significant genetic convergence with GWAS risk loci for SCZ and MDD, placing these cross-lineage states at the epicenter of neurodegenerative and neuropsychiatric pathogenesis.

## Declarations

### Ethics approval and consent to participate

Ethics approval statements for animal experiments: The study adhered to the National Research Council’s Guide for the Care and Use of Laboratory Animals and received approval from the Comments of Animal Experiments and Experimental Animal Welfare Committee of Capital Medical University (approval no. AEEI-2025-968).

### Consent for publication

Not applicable.

### Availability of data and materials

The datasets analysed during the current study are available publicly. The single-cell transcriptome of hippocampus of adult mice is accessible at National Genomics Data Center (NGDC), with GSA: CRA019523 (https://ngdc.cncb.ac.cn/gsa) for raw sequence data and OMIX008044 (https://ngdc.cncb.ac.cn/omix) for processed 10X gene expression matrix; the processed MERFISH spatial transcriptome of adult mice is accessible at https://github.com/AllenInstitute/abc_atlas_access/blob/main/descriptions/MERFISH-C57BL6J-638850.md; the scRNA-seq data for oligodendrocyte lineage trajectory analysis were accessed via the Neuroscience Multi-omic Data Archive (https://assets.nemoarchive.org/dat-61kfys3). Additional datasets, including the 5xFAD model (GSE178304), multiple sclerosis (GSE227781), and Alzheimer’s disease (GSE185553, GSE198323, GSE199243), were retrieved from the NCBI Gene Expression Omnibus (GEO).

### Authors’ contributions

YH conceptualized and supevise the project. YH and YL designed the study. YL and XH conducted data analysis. XH performed the immunofluorescence experiment. YL, XH and YH wrote the manuscript. YN, HW, ZS and JY read and improve the manuscript.

## List of abbreviations

AD: Alzheimer’s disease
APP: Amyloid precursor protein
ARM: Activated response microglia
ASD: Autism spectrum disorder
CNS: Central nervous system
DAM: Disease-associated microglia
DEGs: Differentially expressed genes
DG: Hippocampal dentate gyrus
DPM: Dual-phenotype microglia
GO: Gene Ontology
GSEA: Gene Set Enrichment Analysis
LDAM: Lipid-droplet-accumulating microglia
MDD: Major depressive disorder
mDPM: Myelin-associated dual-phenotype microglia
MFOLs: Myelinating precursor oligodendrocytes
MGnD: Microglial neurodegenerative phenotype
MS: Multiple Sclerosis
MyTE: Myelin-transcript-enriched-microglia
nDPM: Neuron-associated dual-phenotype microglia
OL: Oligodendrocyte
PPI: Protein-protein interaction
RF: Random Forest
scRNA-seq: Single-cell RNA sequencing
snRNA-seq: Single-nucleus RNA sequencing
SCZ: Schizophrenia
TF: Transcription factor
WAM: White matter-associated microglia
WGCNA: Weighted gene co-expression network analysis

## Acknowledgments

We acknowledge the original investigators for making these high-quality datasets publicly available. We thank Liu Mingxia, Ren Siyu, Li Jing, Huang Huanwei, Wang Fei et al.’ participant in work discussion and offering advice.

