## Supplemental Figures S1-S5 for "Single-cell Transcriptomics Analyses Revealed Specialized Microglial Subsets with Oligodendrocyte-like Signatures"

### Supplementary figures

Figure S1. Functional characterization of DPM subtypes.

Figure S2. Computational and morphological characterization of DPM subtypes.

Figure S3. DPM subtype-specific regulon and cell-cell communication.

Figure S4. Cell mapping of DPM subtypes implicated the developmental dynamics.

Figure S5. Integrative Genomics Viewer (IGV) tracks displaying the genomic distribution of sequencing reads for the St18 in mDPMs.

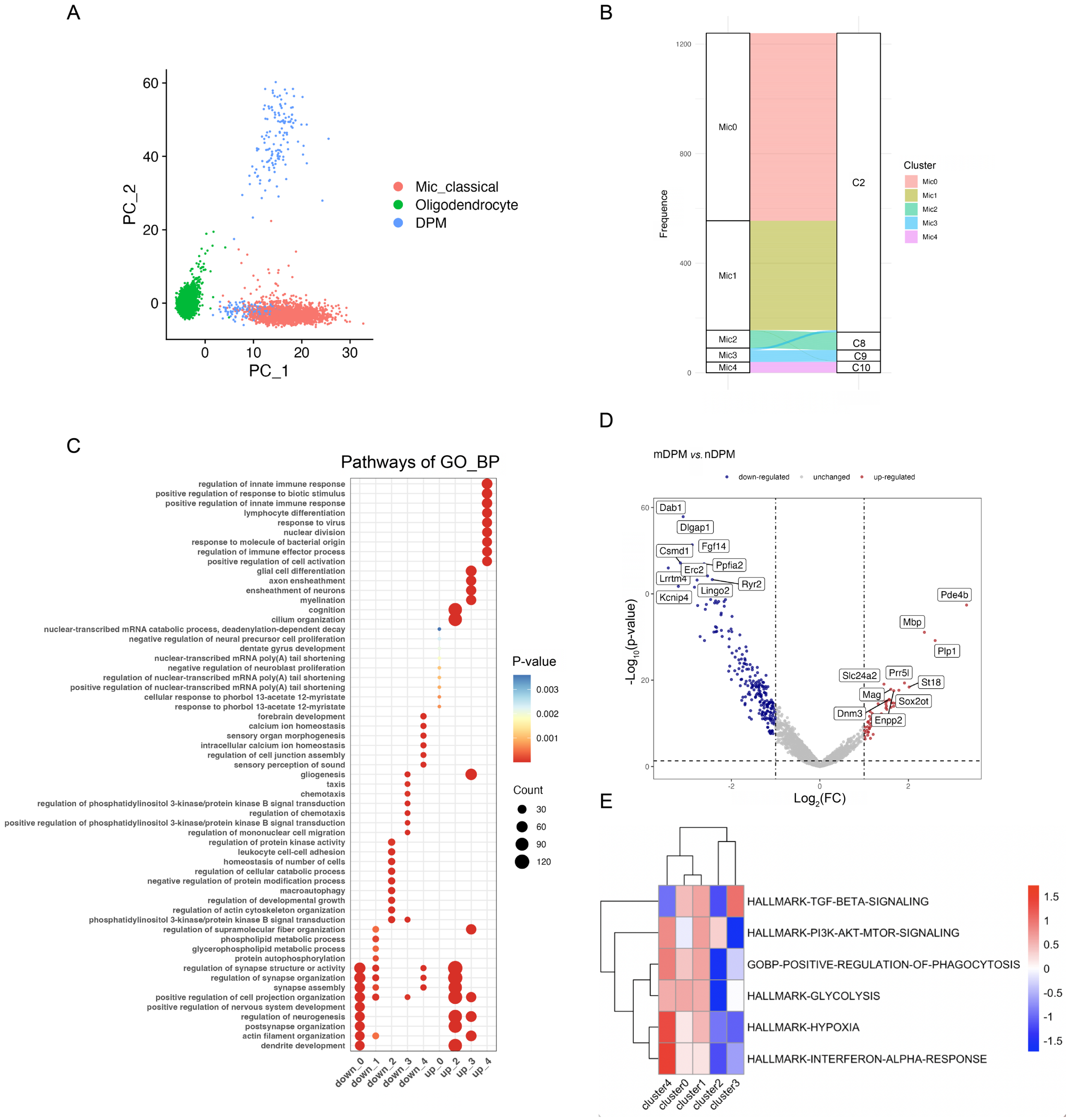


**Figure S1.** Functional characterization of DPM subtypes. **(A)** PCA visualization of the transcriptomic relationships between classical microglia, oligodendrocytes, and the DPM population. **(B)** Sankey diagram illustrating the distribution and cluster transition across different data groupings of the DPM subpopulation. **(C)** Dot plot showing the GO biological process enrichment for upregulated and downregulated genes across clusters 0–4. **(D)** Volcano plot comparing mDPM (Cluster 3) and nDPM (Cluster 2) by limma. Log2 fold change > 1 and *P* < 0.05. **(E)** Heatmap of Gene Set Variation Analysis (GSVA) scores for hallmark pathways and specialized biological processes.

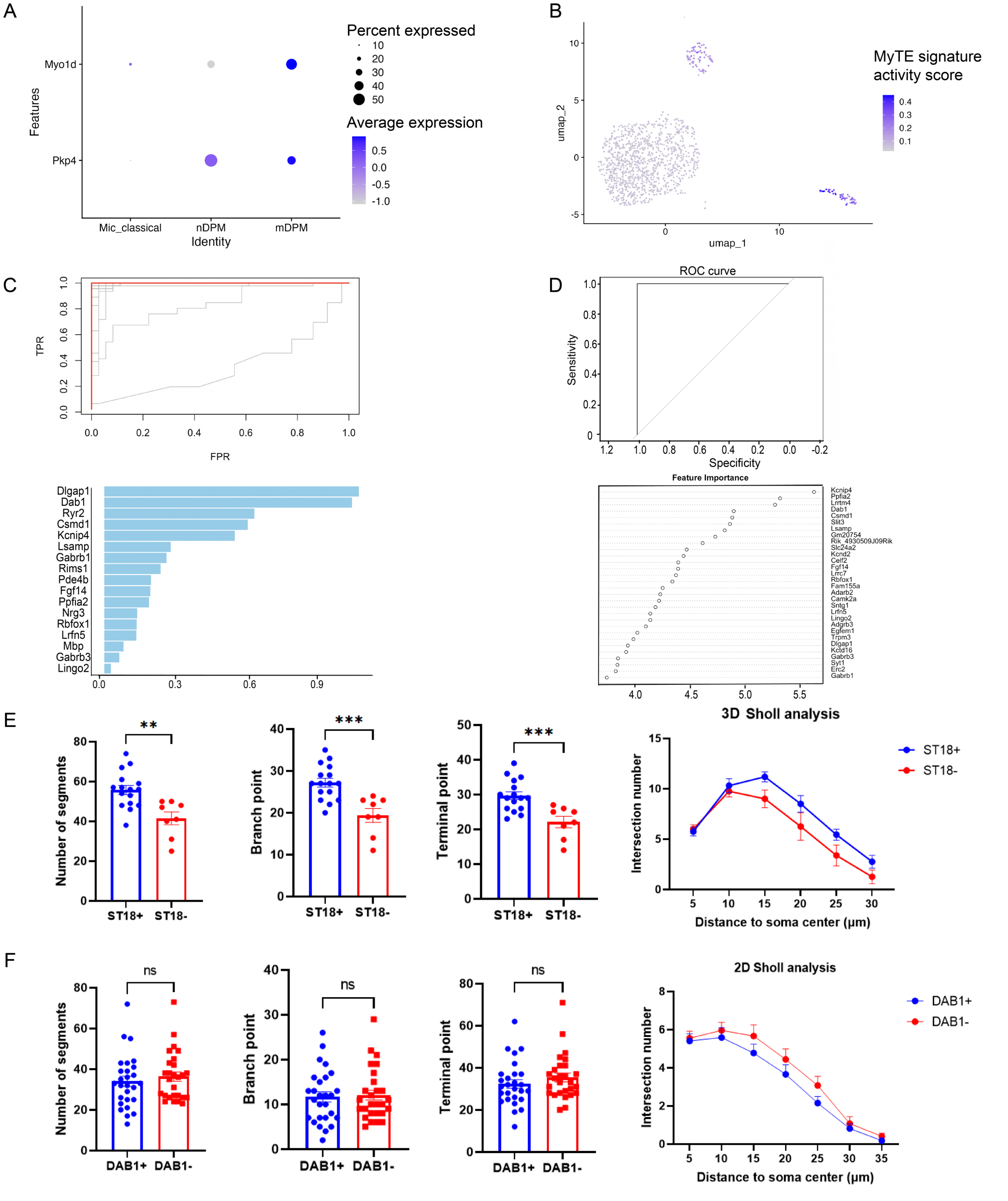


**Figure S2.** Computational and morphological characterization of DPM subtypes. **(A)** Expression of mDPM marker genes in independent datasets. **(B)** AUCell scoring of MyTE signature. **(C-D)** ROC curves and importance rankings of discriminative genes identified by LASSO regression (C) and Random Forest (D) models. **(E)** Quantitative comparison of segments, branching points, and terminal points between ST18+ (n = 16) and ST18- (n = 8) microglia from 3D analysis. 3D Sholl analysis curves (right) showing the number of intersections at increasing distances from the soma center for ST18+ and ST18- microglia. **(F)** (left) Quantitative comparison of morphological parameters derived from 2D reconstruction between DAB1+ (n = 27) and DAB1- (n = 27) microglia. (right) 2D Sholl analysis curves showing the number of intersections at increasing distances from the soma center for DAB1+ and DAB1- microglia. Data are presented as mean ± SEM; unpaired t-test, **P* < 0.05, ***P* < 0.01, ****P* < 0.001.

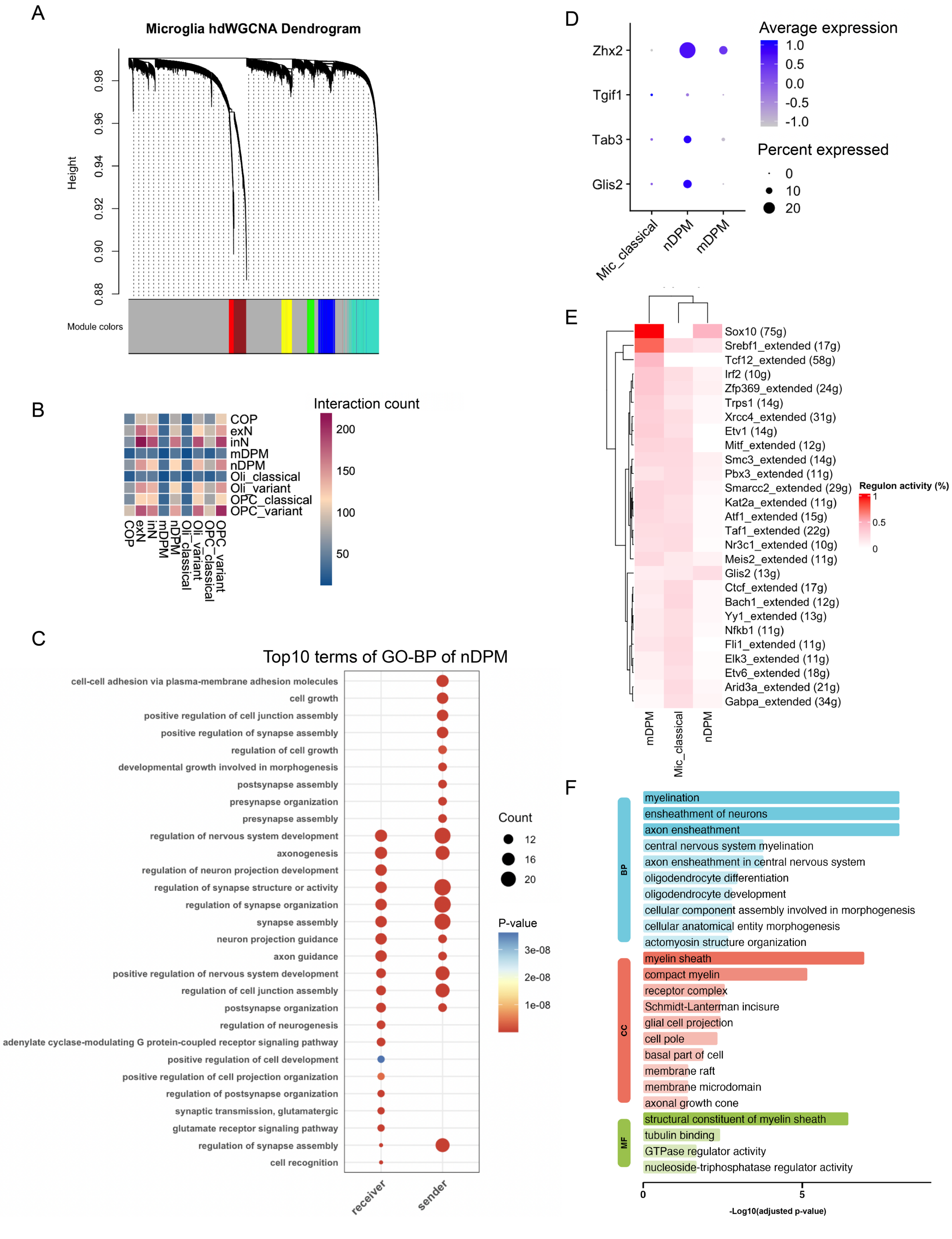


**Figure S3.** DPM subtype-specific regulon and cell-cell communication. **(A)** WGCNA cluster dendrogram. **(B)** CellphoneDB-based intercellular interaction heatmap. **(C)** Specific signaling pathways between nDPM and neurons. **(D)** Specific expression of *Glis2* and its putative target gene. **(E)** Heatmap showing the percentage of cells in each subpopulation with active regulons, based on binarized SCENIC activity thresholds. Only regulons with an activation frequency exceeding 20% in at least one subpopulation (minPerc = 0.2) are shown. **(F)** Functional enrichment of *Sox10* target genes.

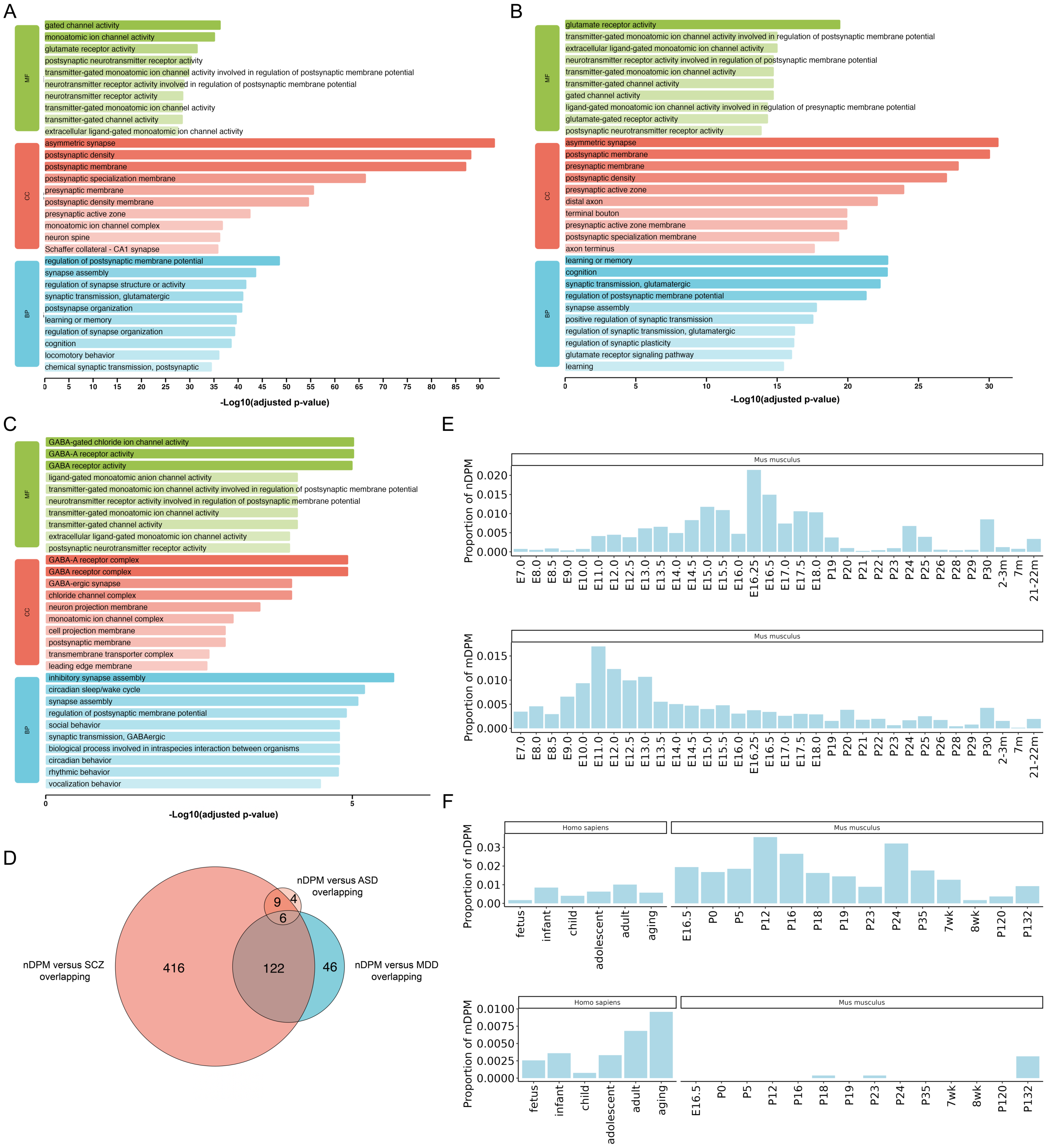

**Figure S4.** Cell mapping of DPM subtypes implicated the developmental dynamics. **(A–C)** Functional pathways for genes intersecting nDPM with SCZ (A), MDD (B), and ASD (C). **(D)** Intersection of nDPM signature DEGs with SCZ, MDD and ASD risk genes. **(E)** DPM proportions across mouse brain development. **(F)** Cross-species hippocampal DPM dynamics.

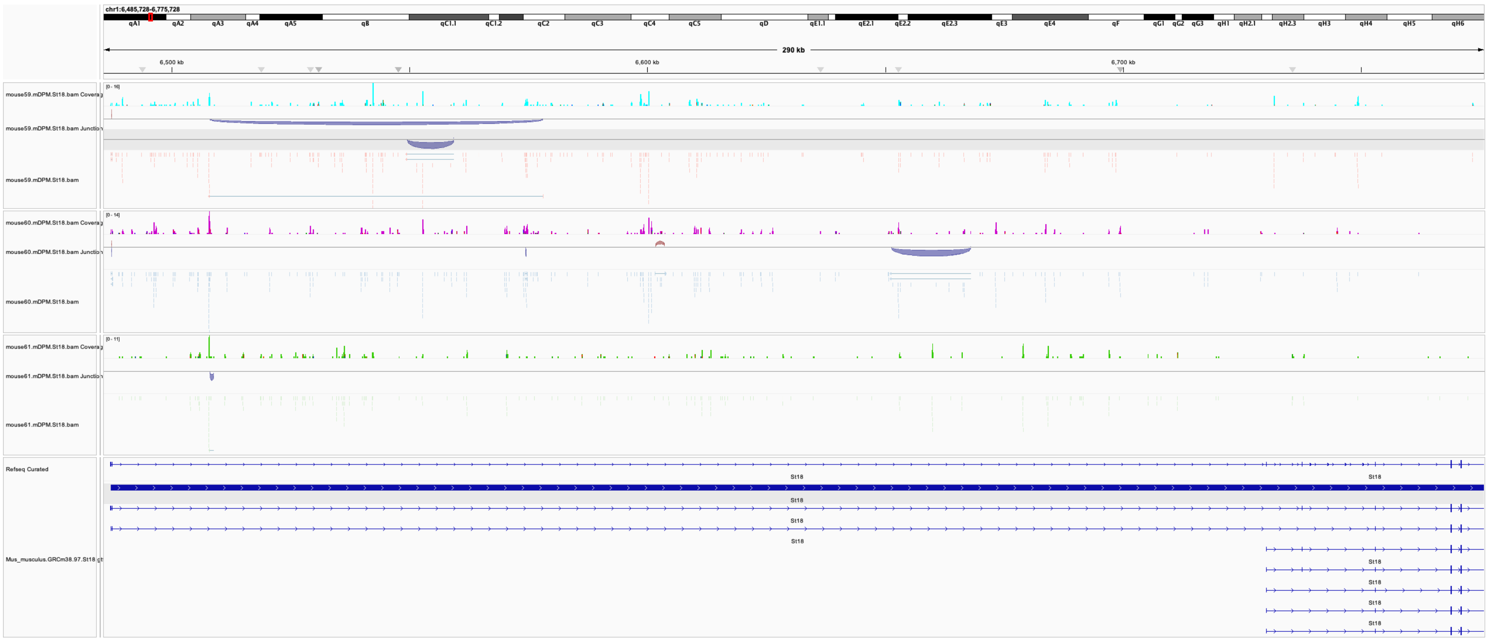


**Figure S5.** Integrative Genomics Viewer (IGV) tracks displaying the genomic distribution of sequencing reads for the *St18* in mDPMs.
